# Identification of biomarkers and candidate regulators for multiple myeloma under the knockout of AURKA

**DOI:** 10.1101/2021.03.21.436324

**Authors:** Hanming Gu, Wei Wang, Gongsheng Yuan

## Abstract

Multiple myeloma (MM) is a plasma cell malignancy that is characterized by the overabundance of monoclonal paraprotein. Aurora kinase A (AURKA) was upregulated in patients with high-risk MM. AURKA inhibitors were used to inhibit MM cell proliferation by inducing cell apoptosis and injury. In our study, we aim to identify biological processes and pathways of MM cells under the knockout of AURKA (AURKA KO) by using a bioinformatics method to elucidate their potential pathogenesis. The gene expression profiles of the GSE163589 dataset were originally produced by using the high-throughput BGISEQ-500 (Homo sapiens). The biological categories and pathways were analyzed by the Kyoto Encyclopedia of Genes and Genomes pathway (KEGG), Gene Ontology (GO), and Reactom enrichment. KEGG and GO results indicated the biological pathways related to the immune responses and cancer activities were mostly affected in the development of MM with AURKA KO. Moreover, we identified several genes including GNG5, UBE2D1, and BUB1B were involved in the regulation of cancer genesis. We further predicted novel regulators that had the ability to affect the progression of MM with AURKA KO based on the L1000fwd analysis. Therefore, this study provides further insights into the mechanism of MM under AURKA inhibitor treatments.

## Introduction

Multiple myeloma (MM) is a common malignancy of terminally differentiated plasma cells^1^. MM cells are originated in the bone marrow, but they also reside in the peripheral blood and other organs^2^. MM accounts for about 1.7% of all malignancies in the US. The incidence of MM is higher in Americans but lower in Asian and Hispanic individuals^1^. MM is characterized by the secretion of monoclonal immunoglobulin proteins that are produced by pathologic plasma cells^3^. The clinical manifestations include monoclonal protein, malignant cells, end-organ damage (bone disease with lytic lesions, anaemia, renal insufficiency, and hypercalcarmia)^4^. MM cells are affected by the bone marrow microenvironment because of the adhesion of MM cells to extracellular-matrix proteins^5^. In addition, binding of MM cells to BM accessory cells induces secretion of cytokines, which further promotes tumor cell activation^6^.

Aurora kinases were found to regulate cell-cycle checkpoints and some related molecules such as cyclins and cyclin-dependent kinases^7^. Aurora kinases localize in the centrosome and play important roles in cell division by regulating chromatid segregation in mitotic cells; moreover, loss of chromatid segregation leads to genetic instability and tumorigenesis^8^. There are three members of the mitotic Aurora kinase family: Aurora-A, Aurora-B and Aurora-C kinases. Aurora-A localizes to centrosomes and associates with spindle microtubules proximal to the spindle poles during mitosis^9^. Enhanced expression of AURKA was reported in several cancers including laryngeal, breast cancers, and MM. Moreover, AURKA associated with cell proliferation, metastasis and chemoresistance^10^. AURKA was recently reported as a crucial target of MM. Upregulation of AURKA is related to centrosome amplification and worse prognosis in MM^11^. Reports showed that inhibition of AURKA gene expression in MM cells by RNAi leads to apoptosis and cell death as well as inhibits G2/M cell-cycle progression in MM cell lines^12^. Thus, regulation of AURKA and related pathways may be a convincing strategy in MM therapy.

In this study, we investigated the effect of the knockout of AURKA (AURKA KO) in MM cells. We identified several DEGs, candidate inhibitors and the relevant biological processes of MM cells with AURKA KO utilizing comprehensive bioinformatics analyses. We performed the functional enrichment analysis and protein-protein interaction for finding significant gene nodes. These key genes and signaling pathways could be critical to therapeutic interventions of MM.

## Materials and Methods

### Data resources

The dataset GSE163589 was downloaded from the GEO database (http://www.ncbi.nlm.nih.gov/geo/). The data was produced by BGISEQ-500 (Homo sapiens), Tianjin Medical University, China. Bulk RNA-Seq analysis was performed using LP-1 that was infected by the virus that packages sgRNA targeting AURKA gene or the non-target control.

### Data acquisition and preprocessing

The dataset GSE163589 containing LP-1 control cells and LP-1 AURKA KO cells samples was conducted by R script^13, 14^. We used a classical t test to identify DEGs with P<.01 and fold change ≥1.5 as being statistically significant.

### Gene functional analysis

Gene ontology (GO) analysis is a useful tool to develop a comprehensive model of biological systems and to provide a system for hierarchically classifying genes^15, 16^. Kyoto Encyclopedia of Genes and Genomes (KEGG) database is a commonly used tool for understanding the biological system, which integrates functional information, biological pathways, and sequence similarity. We performed the GO and KEGG pathway analyses by utilizing the Database for Annotation, Visualization, and Integrated Discovery (DAVID) (http://david.ncifcrf.gov/) and Reactome. P<.05 and gene counts >10 were considered statistically significant.

### Module analysis

The Molecular Complex Detection (MCODE) of Cytoscape software was used to analyze the densely connected regions in protein-protein interaction (PPI) networks^17^. The significant modules and clusters were selected from the constructed PPI network using MCODE. The function and pathway enrichment analyses were performed by using Reactome, and P<.05 was used as the cutoff criterion.

### Reactome and L1000FDW analysis

We performed the Reactom pathway (https://reactome.org/) and L1000FDW (https://maayanlab.cloud/L1000FWD/) to obtain the visualization, interpretation and analysis of potential pathways. P<.05 was considered statistically significant.

## Results

### Identification of DEGs in myeloma cells with AURKA KO

LP-1 cells were infected by the virus that packages sgRNA targeting AURKA gene (Aurora A-KO) or the non-target gene (Control). AURKA KO cells were harvested for gene expression profiling. To gain the insights on LP-1 cells with AURKA KO genes, the modular transcriptional signature of AURKA KO cells was compared to that of the Control cells. A total of 586 genes were identified to be differentially expressed in AURKA KO cells with the threshold of P<0.05. The top 10 up- and down-regulated genes for AURKA KO cells and control samples are list in table 1.

**Table 1.**

| **Entrez gene** | **Gene Symble** | **Fold-change** | **Regulation** |
| --- | --- | --- | --- |
| Top 10 down-regulated DEGs |  |  |  |
| 79805 | VASH2 | -1.825741858 | Down |
| 642597 | AKAIN1 | -1.825741858 | Down |
| 55975 | KLHL7 | -1.81897851 | Down |
| 10142 | AKAP9 | -1.81323663 | Down |
| 26256 | CABYR | -1.81323663 | Down |
| 81567 | TXNDC5 | -1.811197356 | Down |
| 51303 | FKBP11 | -1.810806906 | Down |
| 101928635 | LOC101928635 | -1.80972627 | Down |
| 5701 | PSMC2 | -1.808122644 | Down |
| 284615 | ANKRD34A | -1.807454814 | Down |
| Top 10 up-regulated DEGs |  |  |  |
| 346689 | KLRG2 | 1.825741858 | up |
| 4145 | MATK | 1.825741858 | up |
| 83417 | FCRL4 | 1.825741858 | up |
| 3553 | IL1B | 1.825741858 | up |
| 3362 | HTR6 | 1.825741858 | up |
| 55220 | KLHDC8A | 1.825741858 | up |
| 9087 | TMSB4Y | 1.825741858 | up |
| 1278 | COL1A2 | 1.825741858 | up |
| 10814 | CPLX2 | 1.821415451 | up |
| 5126 | PCSK2 | 1.820356178 | up |

### KEGG analysis of DEGs in myeloma cells with AURKA KO

To identify the biological roles and potential mechanisms of the DEGs from AURKA KO versus Control in LP-1 cells, we performed KEGG pathway enrichment analysis (Supplemental Table S1). KEGG pathway database (http://www.genome.jp/kegg/) is a collection of genes that are highly similar in sequence for understanding the molecular interaction, reaction and relation networks. Our study indicated the top ten enriched KEGG pathways including “Pathways in cancer”, “Proteoglycans in cancer”, “Influenza A”, “Hippo signaling pathway”, “Oxytocin signaling pathway”, “Alzheimer’s disease”, “Cell cycle”, “Lysosome”, “Measles”, and “Leukocyte transendothelial migration” (Figure 1).

**Figure 1.**
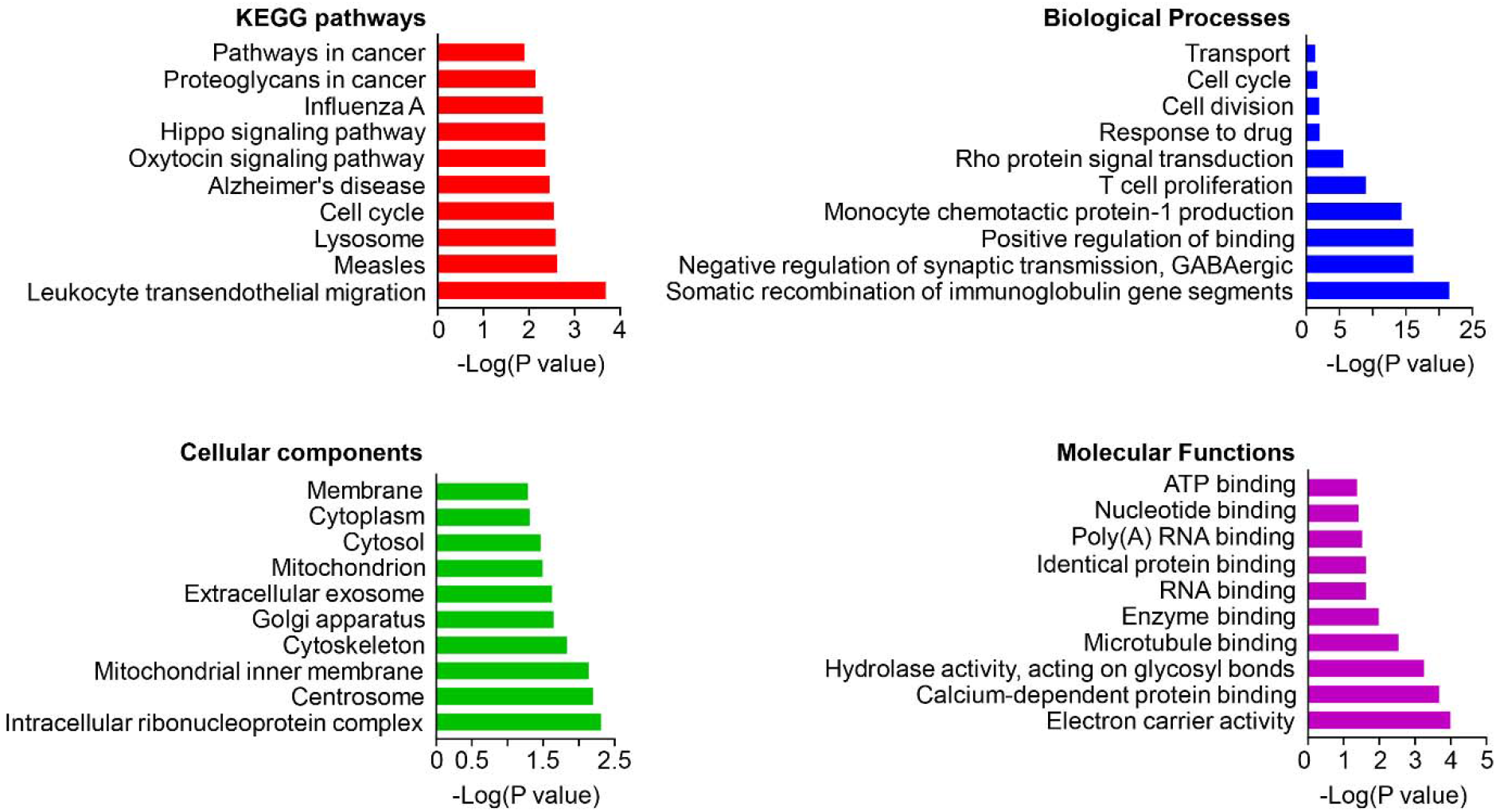
The KEGG pathways, biological process, cellular component, and molecular function terms enriched by the DEGs. DEGs =differentially expressed genes, KEGG = Kyoto Encyclopedia of Genes and Genomes.

### GO analysis of DEGs in myeloma cells with AURKA KO

Gene ontology (GO) analysis is a useful tool for hierarchically classifying genes, which contains cellular components (CC), molecular functions (MF), and biological processes (BP). Here, we identified top ten cellular components including “extracellular exosome”, “cytoplasm”, “membrane”, “cytoskeleton”, “centrosome”, “Golgi apparatus”, “mitochondrion”, “cytosol”, “intracellular ribonucleoprotein complex”, and “mitochondrial inner membrane” (Figure 1). We then identified top ten biological processes: “transport”, “cell cycle”, “cell division”, “response to drug”, “Rho protein signal transduction”, “T cell proliferation”, “positive regulation of monocyte chemotactic protein-1 production”, “negative regulation of synaptic transmission, GABAergic”, “positive regulation of binding”, and “somatic recombination of immunoglobulin gene segments” (Figure 1). We identified top ten molecular functions: “ATP binding”, “nucleotide binding”, “poly(A) RNA binding”, “identical protein binding”, “RNA binding”, “enzyme binding”, “microtubule binding”, “hydrolase activity, acting on glycosyl bonds”, “calcium-dependent protein binding”, and “electron carrier activity” (Figure 1).

### PPI network and Module analysis

We created the PPI networks to identify and analyze the relationships of DGEs at the protein level. The criterion of combined score >0.7 was chosen and the PPI network was constructed by using the 553 nodes and 587 interactions. Among these nodes, the top ten genes with highest scores are shown in Table 2. The top two significant modules of AURKA KO versus Control in LP-1 cells were selected to indicate the functional annotation (Figure 2 and Supplemental Table S2).

**Figure 2.**
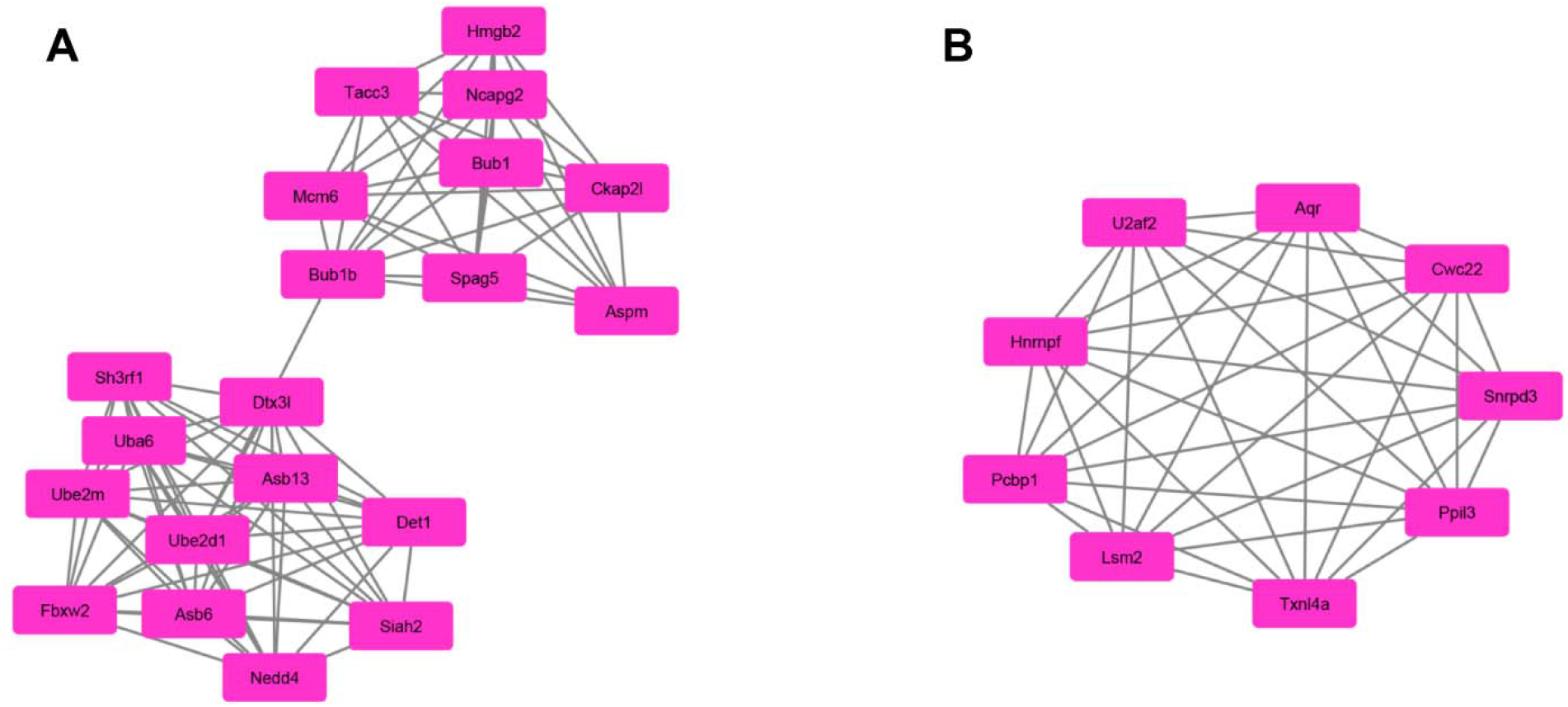
Top two modules (A, module 1 and B, module 2) from the PPI network of MM cells with AURKA KO.

**Table 2.** Top ten genes demonstrated by connectivity degree in the PPI network.

| Gene Symbol | Gene title | Degree |
| --- | --- | --- |
| GNG5 | G protein subunit gamma 5 | 20 |
| UBE2D1 | ubiquitin conjugating enzyme E2 D1 | 17 |
| BUB1B | BUB1 mitotic checkpoint serine/threonine kinase B | 16 |
| ANAPC1 | anaphase promoting complex subunit 1 | 15 |
| BUB1 | BUB1 mitotic checkpoint serine/threonine kinase | 15 |
| KIF18A | kinesin family member 18A | 14 |
| UBE2M | ubiquitin conjugating enzyme E2 M | 14 |
| HLA-DQA2 | major histocompatibility complex, class II, DQ alpha 2 | 13 |
| HLA-DQA1 | major histocompatibility complex, class II, DQ alpha 1 | 13 |
| HLA-DPB1 | major histocompatibility complex, class II, DP beta 1 | 13 |

### Reactome Pathway in myeloma cells with AURKA KO

We identified several signaling pathways by using Reactome Pathway Database (https://reactome.org/). We identified top ten signaling pathways including: “RNA Polymerase I Promoter Opening”, “Amyloid fiber formation”, “DNA methylation”, “ERCC6 (CSB) and EHMT2 (G9a) positively regulate rRNA expression”, “Recognition and association of DNA glycosylase with site containing an affected pyrimidine”, “PRC2 methylates histones and DNA”, “SIRT1 negatively regulates rRNA expression”, “Condensation of Prophase Chromosomes”, “HDACs deacetylate histones”, and “Activated PKN1 stimulates transcription of AR (androgen receptor) regulated genes KLK2 and KLK3” (Supplemental Table S3). We then constructed the reaction map according to the signaling pathways (Figure 3).

**Figure 3.**
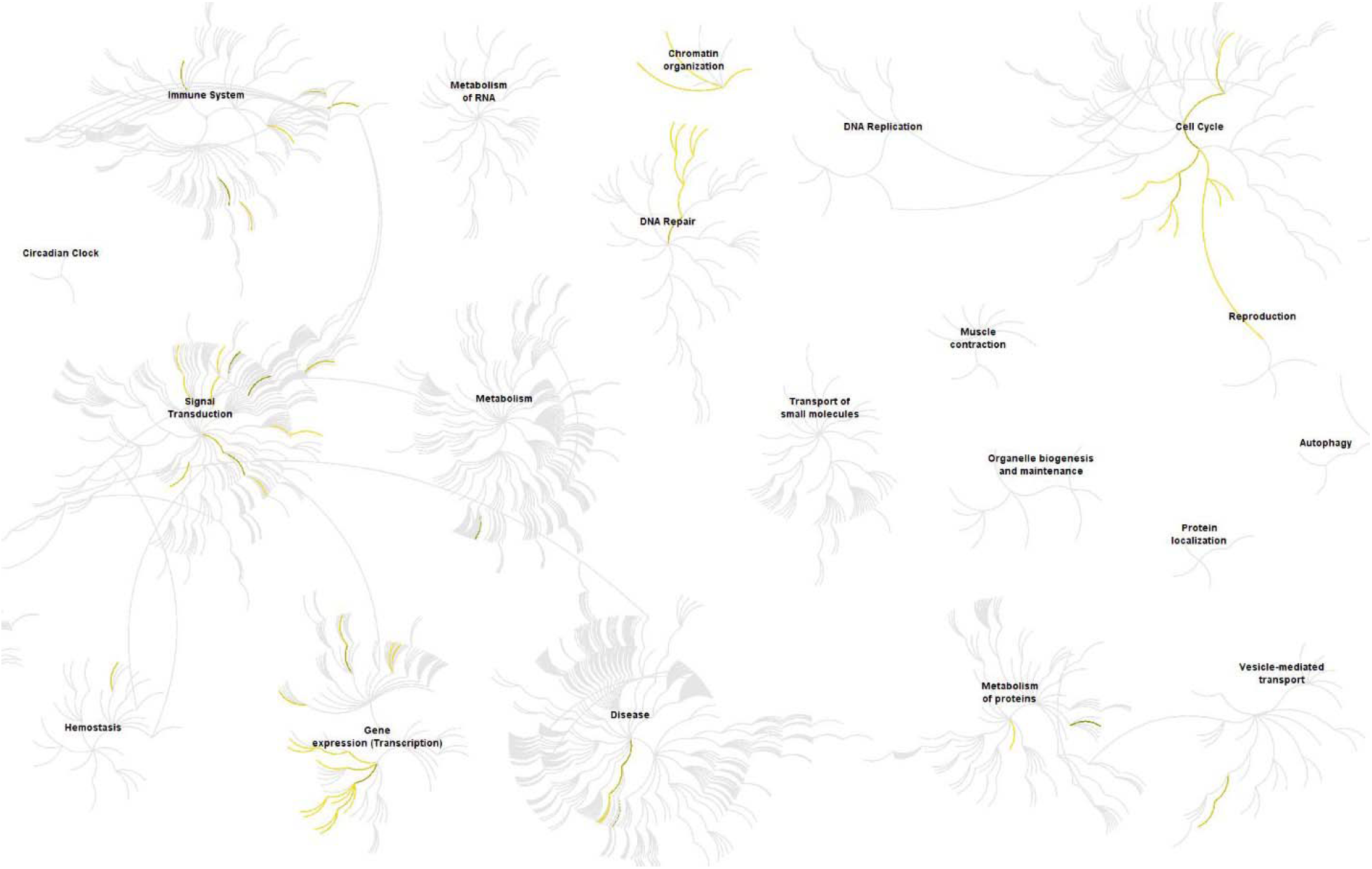
The Reactom pathway visualization map. Input genes are represented by the top significantly changed genes obtained from the GSE163589 dataset (P <0.01). The yellow color represents the most relevant signaling pathways.

**Figure 4.**
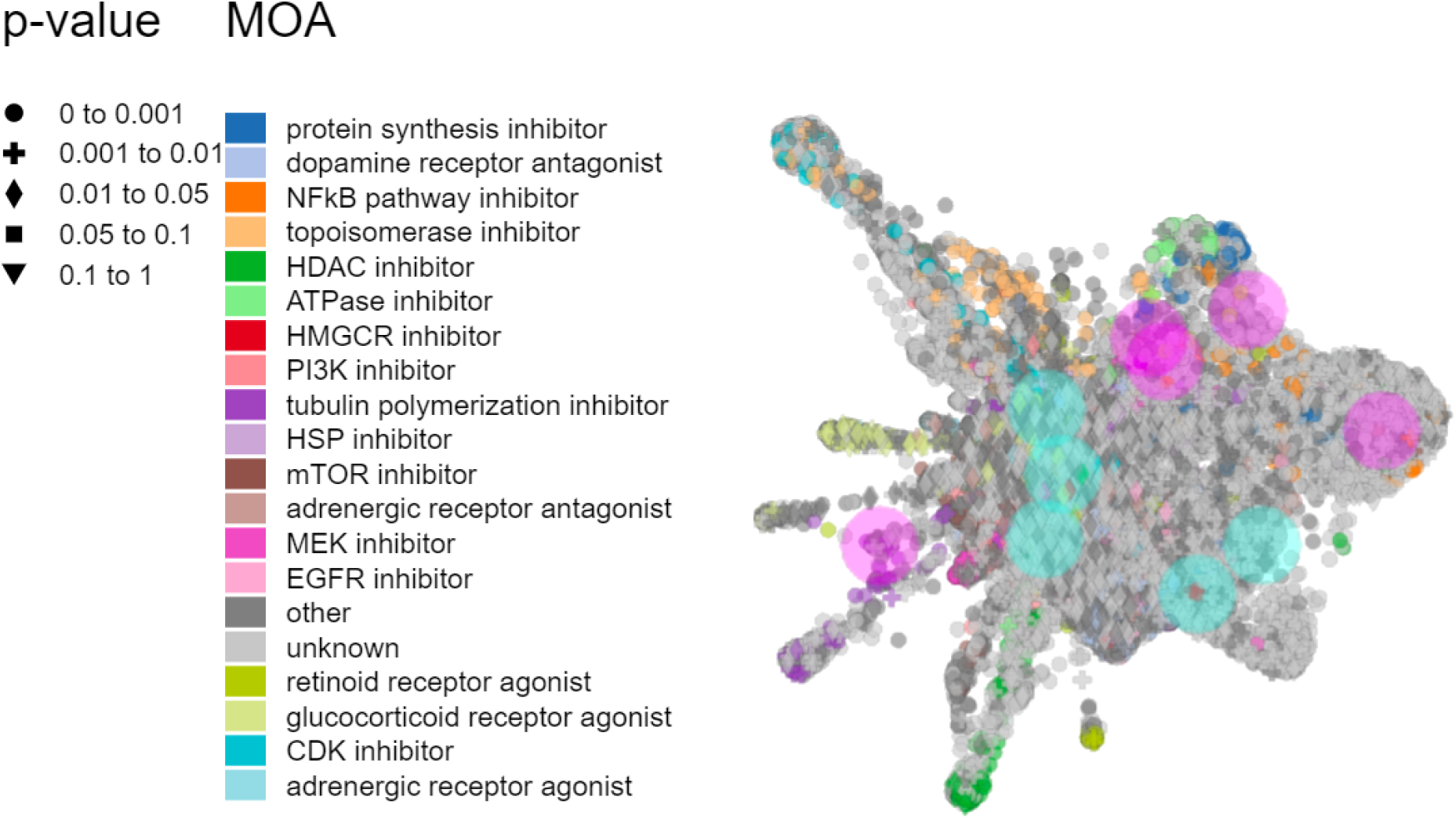
Inhibitor prediction against MM by L1000FDW visualization. Input genes are represented by the significantly changed genes obtained from the GSE163589 dataset. Dots are the Mode of Action (MOA) of the respective drug.

### Potential regulator molecules in AURKA KO cells

To further know the potential regulator molecules, we introduced the L1000FDW system that can predict and analyze the potential bioactive molecules. The system indicated the potential pathways may be blocked. We selected top ten molecules according to the DEGs and the inhibitor map: “BRD-K53501402”, “16,16-dimethylprostaglandin-e2”, “cercosporin”, “BRD-K10484463”, “gemcitabine”, “MD-II-051”, “BRD-A43640821”, “YM-155”, “BRD-K71935468”, and “cymarin” (Supplemental Table S4).

### Discussion

MM is an accumulation of malignant plasma cells in the bone marrow (BM) that leads to bone lesions and immunodeficiency^18^. Inhibition of Aurora-A kinase gene expression in MM cells by RNAi induces apoptosis and cell death. It also affects G2/M cell-cycle in MM cell lines. Inhibitors of pan-Aurora (-A and -B) and Aurora-B kinases are being tested in clinical trials in patients with MM^19^. Moreover, MLN8237 is the first oral inhibitor of Aurora-A kinase, which is currently in early-phase clinical testing in patients with MM^20^. Thus, our study is based on the AURKA KO in MM cell lines, which may provide the gene evidence for clinical trials of AURKA inhibitors.

To understand the effects of AURKA in the MM cells, we analyzed the LP-1 cells with AURKA KO by the Crispa-Cas9 method. By analyzing the DEGs, we selected 10 proteins that may be important during the development of MM after AURKA KO according to the PPI network analysis. G protein, GPCR and RGS play critical roles in pathophysiology^21–23^ and they are involved in numerous diseases such as cancer^24^, bone dysfunction and inflammatory diseases^25–29^. As an important G protein, GNG5 was reported to be increased in gliomas compared with normal samples and it was related to clinicopathologic characteristics^30^. Ube2D1 regulates the ubiquitination of the E3 ubiquitin ligase March-I, which is an independent unfavorable prognostic indicator in lung adenocarcinoma^31^. BUB1B enhances the hepatocellular carcinoma development via triggering the mTORC1 signaling pathway^32^. The ANAPC is an E3 ubiquitin ligase that controls chromosome separation and exit from mitosis in various organisms^33^. BUB1 is reported to enhance the proliferation of liver cancer cells by regulating the SMAD2 phosphorylation^34^. KIF18A enhances the proliferation, invasion and metastasis of HCC cells by promoting the cell cycle signaling pathways and the Akt signaling pathways^35^. UBE2M enhances the cell proliferation through the β-catenin/cyclin D1 in hepatocellular carcinoma^36^. As the novel HLA class II molecules, HLA-DQA2 and HLA-DQB2 genes were expressed in human Langerhans cells^37^. Therefore, these proteins are related to the cancer and immune responses.

KEGG and GO anaylses indicated that immune responses play important roles in the progression of MM cells with AURKA KO. The KEGG and GO anaylses showed the “Leukocyte transendothelial migration”, “T cell proliferation”, and “positive regulation of monocyte chemotactic protein-1 production”, which is associated with the functions of immune cells. Previous reports showed that AURKA can drive early signaling during T-cell activation^38^. Here, our results proved that AURKA KO in MM can significantly affect the expression and activation of immune cells such as T cells. Moreover, we found that Hippo pathways were involved in the progression of MM under the regulation of AURKA. The Hippo signaling pathway is an important mediator of oncogenesis in solid tumors^39^. Recent finding showed that there was a direct link between the Hippo pathway, NF-κB, and circadian clock activity^40^. NF-κB is related to numerous inflammatory diseases such as RA and OA^25, 41, 42^. Circadian clock genes regulate the majority of physiology and pathophysiology processes, such as metabolism, apoptosis, aging and diseases^43–45^. Yap1-mediated NF-κB activity interrupts the circadian oscillation by inhibiting Per1, Per2, and Cry2 levels^40^. Adrian Rivera-Reyes also found the inhibition of the UPR and clock caused by YAP1 support a shift in metabolism toward cancer cell-associated glycolysis and hyper-proliferation^40^. Another interesting finding is that the “Alzheimer’s disease” and “Influenza A” were changed during the AURKA KO in MM, which suggested that the treatment of MM by targeting the AURKA may affect the therapy effects of these two diseases.

Briefly, we identified the potential biomarkers and pathways for MM cells during AURKA KO. Immune dysfunction and cancer pathways are two key processes during the treatment of AURKA KO. Future studies will focus on the administration of potential AURKA inhibitors on clinical trials. This study thus provides further insights into the treatment of MM under inhibiting AURKA, which may facilitate the drug development.

